# *Tityus serrulatus* venom: TsTX-κ beta a neurotoxin with anti-microbial activity and non-hemolytic

**DOI:** 10.1101/2022.10.09.511467

**Authors:** Thiago de Jesus Oliveira, Nayrob Pereira, Katie Cristina Takeuti Riciluca, Soraia Maria do Nascimento, Ursula Castro de Oliveira, Pedro Ismael da Silva

## Abstract

Increase of infections caused by microorganisms resistant to conventional antibiotics is a health problem in Brazil and worldwide. The search for new molecules capable of inhibiting the growth of pathogens is a challenge for researchers, who find in venoms a rich source of biomolecules, including antimicrobial peptides (AMPs). The Brazilian scorpion, *Tityus serrulatus*, is one of the species that cause serious accidents; its venom is rich in neurotoxins that have been well characterized, highlighting their activities on channels (especially sodium and potassium). In this study we identifified and characterized one AMPs in *T. serrulatus* venom. After milking, the venom was fractioned by high performance liquid chromatography and the fractions were tested by liquid growth inhibition assay, the minimum inhibitory concentration (MIC) against *Escherichia coli, Micrococcus luteus, Candida albicans* and *Aspergillus niger*. The fraction identified with antimicrobial activity was analyzed by electrophoresis and mass spectrometry and this AMP (with molecular mass 6.882 kDa) has a similar amino acid sequence to TsTX-κ beta, a neurotoxin that acts on ion channels. The TsTX-κ beta in this study was identified by *de novo* sequencing. This peptide showed activity against all microorganisms tested. At high concentrations, this peptide, showed hemolytic activity against human erythrocytes. This is a new function described for this peptide, the identification of antimicrobial activity in a neurotoxin already known.

**Key Contribution:** Multifunction: antimicrobial and hemolytic activity associated to TsTX-κ beta, a neurotoxin that acts on potassium channels.

## 1. Introduction

In Brazil, the scorpion *Tityus serrulatus* (family Buthidae), is found in different habitats, especially in the south-eastern region of the country (Cupo *et al*. 1994). This wide distribution is justified by different reasons such as good adaptation in urban and rural environments, great food supply, reproduction by parthenogenesis, an effective immune system, efficient apparatus for capturing prey and an excellent defense against predators (Lourenco and Cloudsley-Thompson 1999). These last two advantages are mainly due to a complex cocktail of biologically active molecules present in the poison (Rodríguez de la Vega *et al*. 2004). The severity of the poisoning caused by *T. serrulatus* is explained by the complexity of its venom, rich in insoluble mucus, bioactive amines, enzymes (metalloproteases, metallopeptidases, hyaluronidases, serine proteases, and peptidases), bradykinin-potentiating peptides, allergenic proteins, ion channels toxins (potassium (K+) and sodium (N+)), and bioactive peptides (antimicrobial peptides (AMPs) and antitumoral peptides) (Bordon *et al*. 2021). The identification of peptides presents from the venom of scorpion has been increasingly explored with the help of “omics tools” (genomics, transcriptomics, metabolomics, and proteomics) (Alvarenga *et al*. 2012, de Oliveira *et al*. 2018).

Among the bioactive molecules present in the scorpion venom, peptides are promising molecules for wide applicability in different pathological treatments. It is possible to classify the peptides into three large classical groups, according to its structural characteristics:

1. peptides with an over representation of certain amino acid residues, such as histidine, proline, or tryptophan. Serrulin is an AMP representative of this group, a peptide glycine-rich isolated from the hemolymph of *T. serrulatus* showing antimicrobial activity against Gram-positive and Gram-negative bacterias, fungus and yeast (Oliveira *et al*. 2019);
2. non-disulfide-bridged peptides (NDBPs), such as linear peptides, examples of NDBPs are pandinin-2 isolated from the *Pandinus imperator* venom (Corzo *et al*. 2001), bioactive molecules that have been synthesized with amino acid sequences obtained by transcripts from the scorpions *Tityus obscurus, Hadrurus gertschi* and *Opistophthalmus cayaporum* venom gland (Guilhelmelli *et al*. 2016) and the linear amidated peptides TsAP-1 and TsAP-2 and their respective analogs (TsAP-S1 and TsAP-S2) were characterized for their antimicrobial activity against *Staphylococcus aureus* ( NCTC 10788), *Candida albicans* (NCPF 1467) and *Escherichia coli* (NCTC 10418), hemolytic activity against horse erythrocytes and antitumor activity against squamous carcinoma cell line (NCI-H157), human lung adenocarcinoma cell line (NCI-H838), and androgen-independent prostate adenocarcinoma (PC-3) (Guo *et al*. 2013).
3. disulfide-bridged peptides (DBPs) they have three or four disulfide bridges and, in most cases, acts on ion channels (Na^+^ or K^+^) (Cerni *et al*. 2014, Peigneur *et al*. 2015). TsTX-κ beta, a neurotoxin that acts on potassium channels (K^+^) is also known by other names such as: beta-κTx 1, Tityustoxin κ-beta, TsTX κ beta, TsTX-κ beta, TsTXκbeta, Ts8, Tsκ2 and scorpine-like (Rogowski *et al*. 1994, Pucca *et al*. 2016). Ts1, as well as Ts8 it is also a beta-neurotoxin, in addition Ts1 shows antifungal activity and no hemolytic activity, this molecule has 3 disulfide bonds and shares similarity with molecules described in other arthropods: Drosomycin (*Drosophila melanogaster*) and Bactridin-1 and 2 molecules from the venom of scorpion *Tityus discrepans* (Santussi *et al*. 2017, Matthews *et al*. 2015, Díaz *et al*. 2009).

All of these studies carried out with the molecules mentioned above are of great importance for the search for new pharmacological targets. The number of microorganisms resistant to antibiotics currently available is growing and the search for candidates for new drugs is becoming increasingly necessary (Linden 2002, Lambert and Lambert 2003). Some of the advantages of using AMPs from a natural source, such as scorpion venom, are: the possibility of finding molecules with a preferential interaction, with specific microbiological targets, this prevents them from interacting with structures that characterize them as toxic to mammalian cells (Peters *et al*. 2010); broad spectrum of activity, being able to act on targets common to many microorganisms or specific to each species; speed in action to contain microbial growth (Shai 2002), mainly compared to the time required for the production of antibodies in the individual and finally; the rare development of microbial resistance to AMP (Peschel and Sahl 2006).

This study reveals new roles for an already known neurotoxin, TsTX-κ beta, present in the venom of *T. serrulatus*, this molecule showed antimicrobial activity against different microorganisms although it also presents hemolytic activity in high doses when tested against human erythrocytes. This work contributes to the characterization of a multifunctional molecule that could become an important pharmacological tool or a future pharmaceutical product.

## 2. Results and Discussion

### 2.1 Reverse-phase High-performance Liquid Chromatography (RP-HPLC)

After milking the *Tityus serrulatus* venom by electrostimulation, the crude venom was centrifuged at 14000 rpm for 10 minutes, collected the supernatant and subjected to high performance liquid chromatography by reverse phase (RP-HPLC) using an analytical column C18. We used 4 mg/mL of venom and the forty-four peaks collected manually analyzed for their activity in inhibiting antimicrobial growth in liquid medium. One of the fractions, denominated fraction 31 (Fig. 1A), showed antimicrobial activity against the following microorganisms: *Escherichia coli* SBS 363 (Gram-negative bacteria), *Micrococcus luteus* A270 (Gram-positive bacteria), *Candida albicans* MDM 8 (yeast) and *Aspergillus niger* (filamentous fungi). To identify the homogeneity of the sample, we submitted fraction 31 to one more fractionation step, using an analytical column C18, however in a different linear acetonitrile (ACN) gradient (20%-50%) and performed the tests again to identify the antimicrobial fraction (inset Fig. 1B). Only one fraction showed antimicrobial activity in the tested conditions.

**Figure 1.**
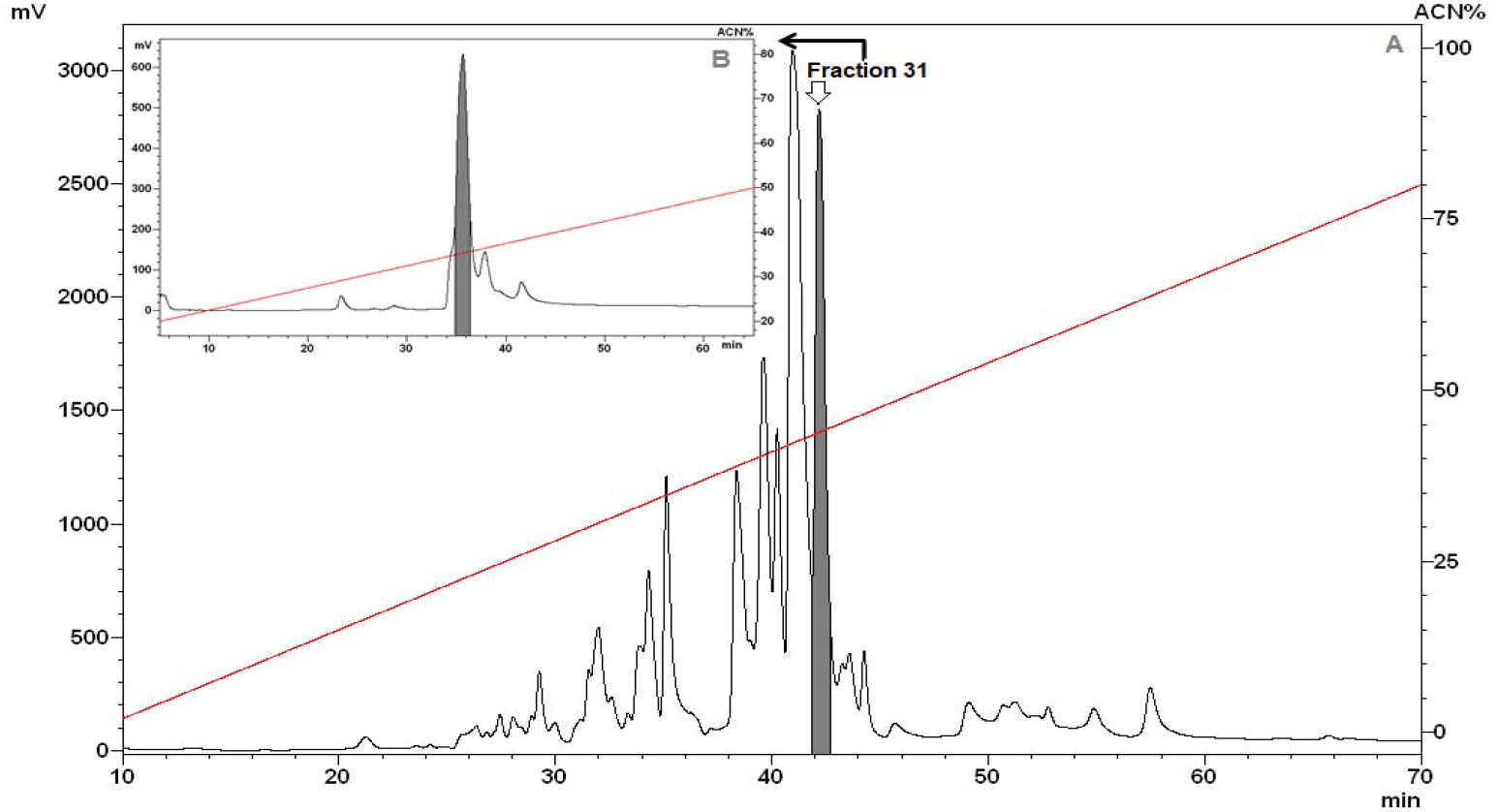
Purification of TsTX-κ beta by reverse-phase high-performance liquid chromatography (RP-HPLC) of *Tityus serrulatus* venom. (A) Fractions eluted with a linear gradient from 2% to 80% acidified ACN (red line) in 0.05% TFA for 60 min (1 ml/min.). The all fractions collected manually were tested on the inhibition of microbial growth in a liquid medium. The grey peak showed antimicrobial activity against the microorganisms: *Escherichia coli* SBS 363, *Microccocus luteus* A270, *Candida albicans* MDM 8 and *Aspergillus niger*. (B) The fraction 31 (white arrow) was submitted to another RP-HPLC using a Jupiter^®^ C18 analytic-column, with a linear gradient from 20% to 50% acidified ACN (red line) in 0.05% TFA, for 60 min at flow rate 1 ml/min. The grey peak TsTX-κ beta showed antimicrobial activity.

### 2.2 Antimicrobial Activity and Minimum Inhibitory Concentration (MIC)

Antimicrobial activity of TsTX-κ beta was screened using only one Gram-positive bacteria (*Micrococcus luteus)*, one Gram-negative bacteria *(Escherichia coli)*, one yeast (*Candida albicans)* and one filamentous fungi (*Aspergillus niger)*, once we had the native TsTX-κ beta, and the minimum inhibitory concentration (MIC) was defined by concentrations ranging from 0.04 to 46.21 μM (Table 01). MICs are expressed as the (a) and (b) interval of concentrations where (a) is the highest concentration tested at which the microorganisms are growing and (b) is the lowest concentration that causes 100% growth inhibition (Ehret-Sabatier *et al*. 1996).

**Table 1.** Spectrum of antimicrobial activity of TsTX-κ beta. The minimal inhibitory concentrations (MICs) of the native peptide was determined by liquid growth inhibition. MICs are expressed as the (a) and (b) interval of concentrations where (a) is the highest concentration tested at which the microorganisms are growing and (b) is the lowest concentration that causes 100% growth inhibition.

| Microorganism | MIC (μM (μg/mL))<sup>1</sup> |
| --- | --- |
| **Gram-negative bacteria** |  |
| *Escherichia coli* SBS 363 | 0.73 – 1.45 μM / (4.96 – 9.93 μg/mL) |
| **Gram-positive bacteria** |  |
| *Micrococcus luteus* A270 | 5.78 – 11.55 μM/ (39.75 – 79.50 μg/mL) |
| **Fungus** |  |
| *Aspergillus niger* | 23.10 – 46.21 μM/ (159 – 318 μg/mL) |
| **Yeast** |  |
| *Candida albicans* MDM 8 | 23.10 – 46.21 μM/ (159 – 318 μg/mL) |
| The highest concentration tested was 46.21 μM/318 μg/mL. |  |

The antimicrobial activity of TsTX-κ beta presents a broad spectrum against different types of microorganisms. The molecule showed a microbial activity against all microorganisms tested, with the concentration for *Escherichia coli* SBS 363 being the lowest *0*.*73 – 1*.*45* μM followed by *Micrococcus luteus 5*.*78 – 11*.*55* μM and filamentous fungus *Aspergillus niger* and yeast *Candida albicans* MDM 8 the MIC of TsTX-κ beta was the same at *23*.*10 – 46*.*21* μM.

When comparing the MICs of TsTX-κ beta with Serrulin, a glycine-rich antimicrobial peptide isolated from the *T. serrulatus* hemolymph, we can observe that TsTX-κ beta has lower concentrations of action only for *Escherichia coli* SBS 363 (*0*.*73 – 1*.*45* μM). (Oliveira *et al*. 2019).

Antimicrobial activity in the venom of the scorpion *T. serrulatus* has already been observed before with TsAP-1 and TsAp-2 identified and analyzed with other approaches, which involve cloning techniques of antimicrobial peptides obtained from cDNA encoded precursors closely related to species *Tityus costatus* and the synthesis of these peptides and their analogs, in a solid phase (Guo *et al*. 2013). The other study involves the different methodology for obtaining Ts1, an antifungal peptide described in the *T. serrulatus* venom (Santussi *et al*. 2017).

### 2.3 Hemolytic Assay

One of the premises for the systematic use of an antimicrobial peptide is that it has low toxicity against human erythrocytes (Oddo *et al*. 2017). The hemolytic assay was performed using human red blood (erythrocytes) of type A+. A serial dilution from the sample at the range of 0.04 to 46.21 μM indicated that in the concentrations above 2.89 μM TsTX-κ beta was toxic to human erythrocytes (Fig. 2) different from TS1 that didn’t show hemolysis at this concentration (*Santussi et al. 2017*).
Is interesting to compare TsTX-κ beta hemolytic activity with the minimum inhibitory concentrations (MICs) obtained from this fraction against the microorganisms previously tested. The values of TsTX-κ beta MICs indicate that the value for *Escherichia coli* SBS363 (*1*.*45* μM) are below the concentration at which a molecule has hemolytic activity (2.89 μM). This highlights the importance of TsTX-κ beta concentrations, which hypothetically could be used as an antibiotic instead antifungals.
Some species of scorpions can show hemolytic activity in their venom, such as that of the species *Opisthancanthus elatus* (Gervais, 1844) from Colombia, this hemolytic activity is associated with action of some toxins, such as enzymes with phospholipase A2 activity performed by serine-proteases (Estrada-Gómez *et al*. 2015). The TsTX-κ beta, is a toxin that belongs to the β-KTx family, these toxins are peptides divided into two domains, N-terminal helical domain (NHD) and C-terminal CSαβ domain (CCD). These sites are related to cytotoxic activities in the NHD and neurotoxic activities in the CCD domains (Pucca *et al*. 2016).
It is important to emphasize that studies carried out with TsTX-κ beta have already been evaluated by other authors regarding their hemolytic activity, however some points must be noted; (1) the concentration used in the evaluation of this activity by Pucca et al. was, 4 μg/100μL (40 μg/mL), while in our work the toxin had hemolytic activity in concentrations above 2.89 μM (19.87 μg/mL). (2) We carried out the tests with human blood (A+), comparing with the work mentioned, fresh mouse blood was used (Pucca *et al*. 2016). Our results of hemolytic activity may suggest that there is a specific target in human cells that explains the toxicity of TsTX-κ beta in concentrations above 2.89 μM.

### 2.4 Sodium Dodecyl Sulfate-Polyacrylamide Gel Electrophoresis (SDS-Page)

The analysis of TsTX-κ beta by Sodium Dodecyl Sulfate-Polyacrylamide Gel Electrophoresis (SDS-PAGE) revealed a band below 10 kDa (Fig. 3). In the process of characterizing the action of TsTX-κ beta in Kv4.2 channel, its sequence of amino acids was defined by N-terminal sequence verified by automated Edman degradation: “KLVALIPNDQLRSILKAVVHKVAKTQFGCPAYEGYCNDHCNDIERKDGECHGFKCKCAKD”, with a molecular weight of 6712 Da corresponding to this toxin (Pucca *et al*. 2016). The result of the mass below 10 kDa observed in SDS-PAGE in addiction to antimicrobial activity confirms our suggestion of an antimicrobial peptide (AMP). No other bands were observed in the sample using this technique. The mass of an antimicrobial peptide gives us information on how many amino acids approximately this molecule has, but it is not possible to obtain the amino acids sequence. Regarding the mass of an antimicrobial molecule, it is possible to find small sequences of amino acids or agents with low molecular weight. An example is 1,4-Benzoquine, a microbial agent isolated from the venom of the scorpion *Diplocentrus melici, molecular mass of 168*. *15 Da and being effective against Staphylococcus aureus* (Carcamo-Noriega *et al*. 2019). And we can find antimicrobial activity associated with larger molecular weight molecules, such as LaIT2 described in the scorpion *Liocheles australasiae*, a molecular mass of 6,628.2 Da which is effective against Gram-negative bacteria and showed insecticidal activity (Juichi *et al*. 2018).

**Figure 2.**
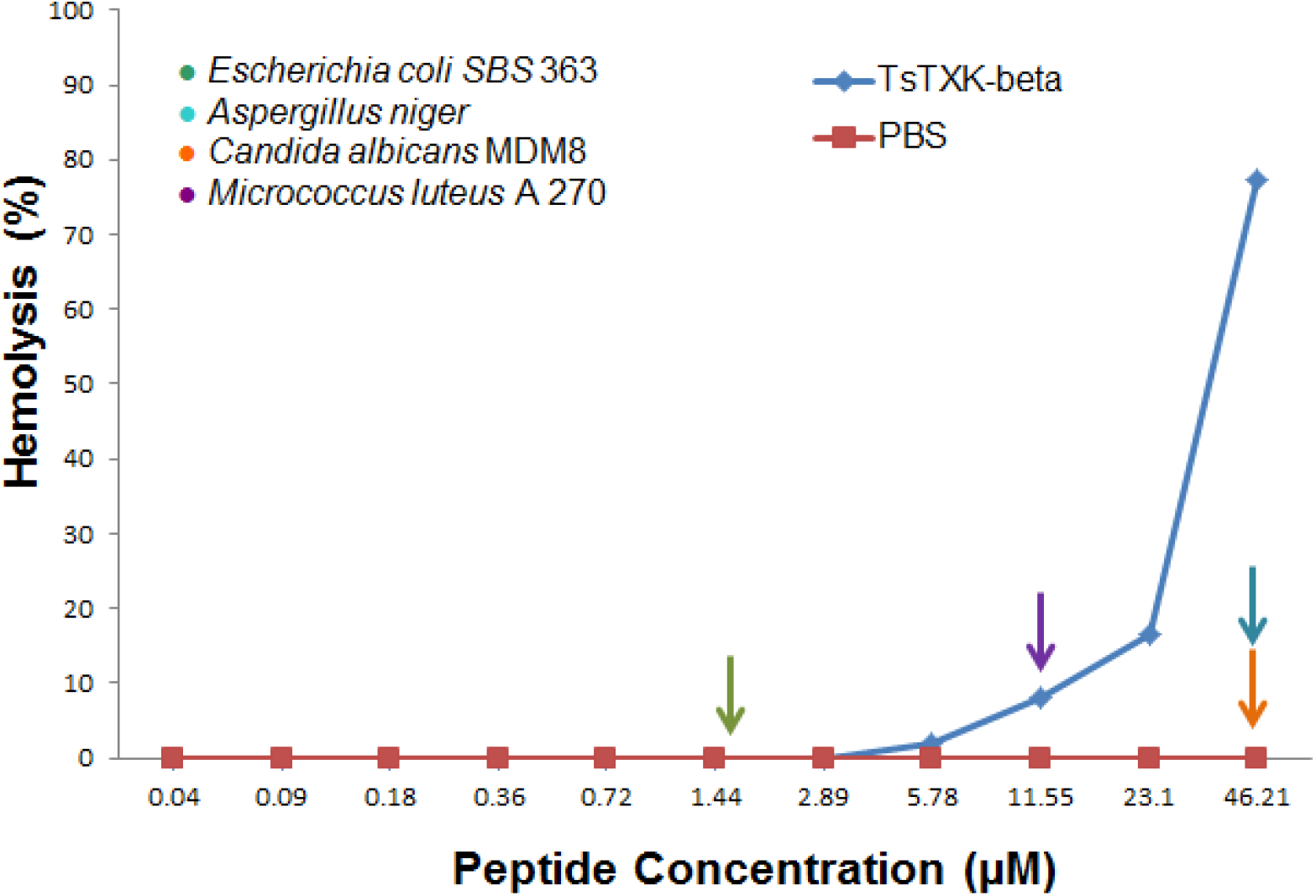
Hemolytic action of TsTX-κ beta against human erythrocytes. The hemolytic activity of TsTX-κ beta was evaluated by means of a serial dilution, obtaining concentrations of 0.04 - 46.21 μM of the sample in the presence of human erythrocytes type A+ (blue line). Hemolysis values are represented in percentage (%), with 100% of the values referring to the positive control for hemolysis (Triton 0.1%). As a negative control we used PBS (phosphate-buffered saline). The arrows in the figure indicate the minimum inhibitory concentrations (MICs) of the TsTX-κ beta, used in the *microbial growth inhibition test in liquid medium, where, for Escherichia coli SBS 363 was 1*.*45 μM (green arrow), Micrococcus luteus* A 270 11.55 μM (purple arrow), *Aspergillus niger* 46.21 μM (blue arrow) and *Candida albicans* MDM 8 46.21 μM (orange arrow).

**Figure 3.**
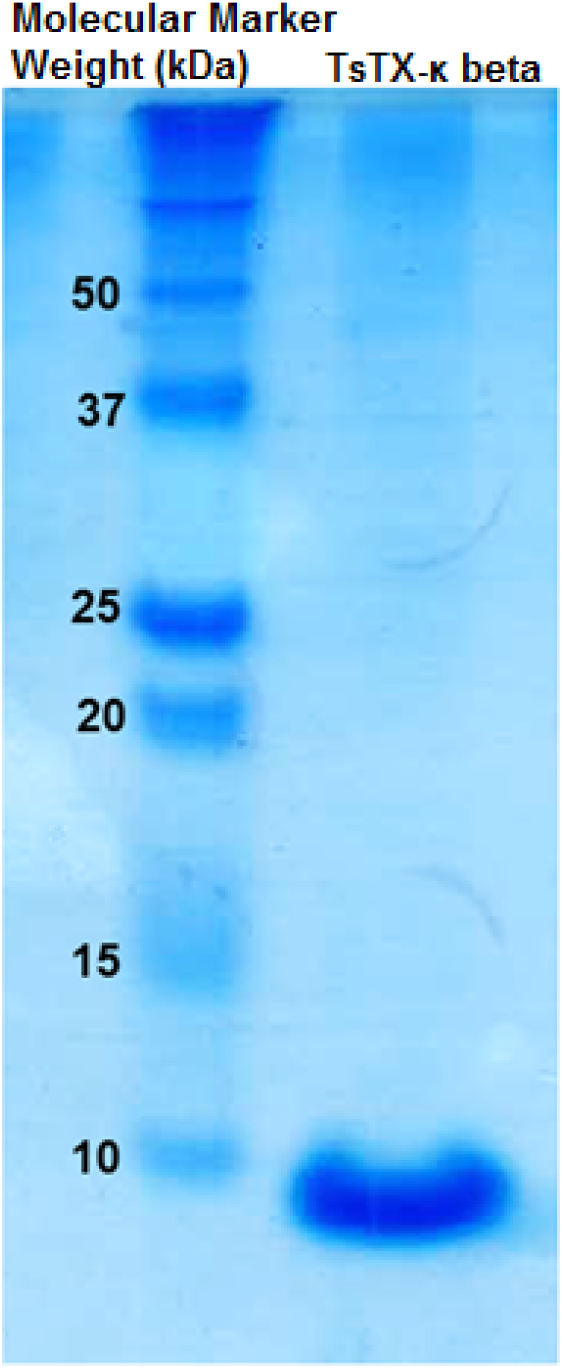
TsTX-κ beta electrophoretic analysis. Column 1: Molecular weight marker (MW) expressed in kDa. Column 2: 50 μg of TsTX-κ beta in a non-reducing condition, was applied to a Sodium Dodecyl Sulfate Polyacrylamide Gel Electrophoresis 20% (20% SDS-PAGE) and stained with Coomassie Brilliant Blue (Bio-Rad, Hercules, CA, USA). When comparing the MWs and the sample, it is possible to observe a band close to 10 kDa., conforming that the biological sample is a peptide.

It is possible to relate the action of antimicrobial peptides on scorpion venoms with different masses and structures. The presence of disulfide bridges classify some antimicrobial peptides as DBS, or for some authors “short scorpion toxins”. These toxins have 23-64 amino acid residues and can form three to four disulfide bridges in different conformations, usually acting on potassium channels (Rodríguez de la Vega and Possani 2004). However, in the scorpion venom it is also possible to find long peptides, these peptides are part of the superfamily of “long scorpion toxins”, they are formed by sequences of 55-76 amino acid residues, generally forming four disulfide bridges (Ortiz et al. 2015). It is possible to highlight molecules that act in sodium channels, for example, the toxin from *T. serrulatus* venom, Ts2 (β-toxin) acts on sodium (Na) channels with different voltages (V): Na(V)1.2, Na(V)1.3, Na(V)1.5, Na(V)1.6 and Na(V)1.7, but does not affect Na(V)1.4, Na(V)1.8 or DmNa(V)1, the latter is an isoform of these channels in insects (Cologna *et al*. 2012). The mass observed in the SDS-Page gel gives us a reference of what is the molecular mass of the sample we are analyzing, however it is not possible to associate the antimicrobial activity of this toxin solely with this information.

### 2.5 Mass Spectrometry and Bioinformatics Analyses

#### 2.5.1 Mass Analysis

Interpretation of the data obtained by mass spectrometry was performed by bioinformatics tools. In order to analyze the structural characteristics of TsTX-κ beta by mass spectrometry, fragments of the molecule obtained by digestion in solution and digestion in gel were used. The raw files were analyzed by the MagTran^®^ tool, where it was possible to identify a m/z of 6881.9 Da (Figure 4), this mass is very close to the mass related to TsTX-κ beta (6712 Da. obtained by Edman degradation) (UniProt 2019). The analysis were carried out using the sample after digestion, it could explain the increase of molecular mass.

**Figure 4.**
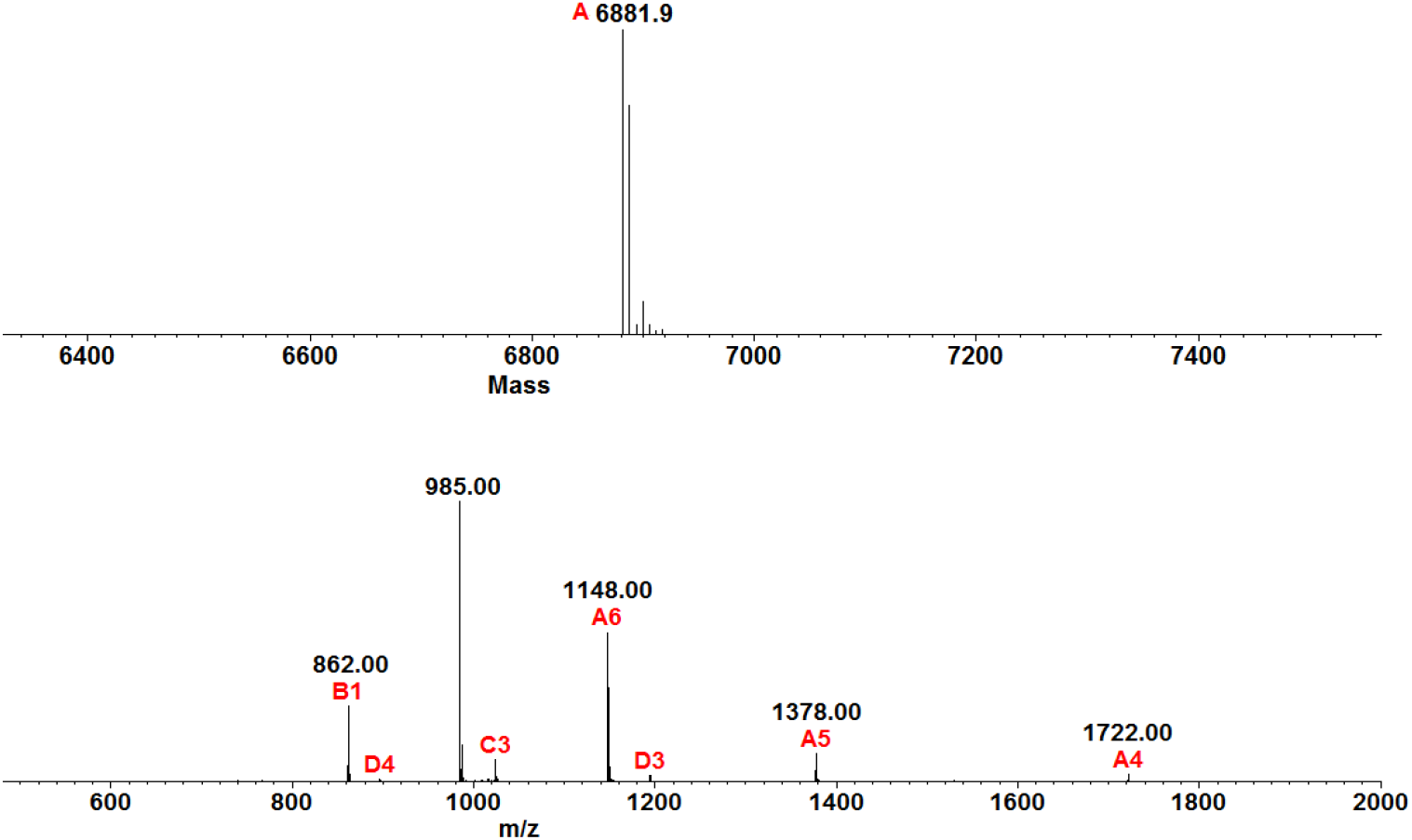
TsTX-κ beta spectrum. Mass spectrometry analyses (LC-MS/MS, coupled with a LTQ-Orbitrap Velos) of TsTX-κ beta revealed an m/z of 6881.9 Da. Ions were submitted to MagTran^®^ for mass analysis and identification.

This mass identified by mass spectrometry, corroborates with the mass of the band observed in the 20% SDS-PAGE (Figure 3), confirming our suspicions that we are analyzing the same sample using different approaches. This mass characterizes TsTX-κ beta as a “long scorpion toxins” (55-77 aa).

#### 5.1.2 Analysis of the TSTxk-beta structure

It was identified by Masslynx^®^ program, the amino acid sequence of one fragment after the method of sample digestion: “LVALIPNDQLR”, referring to our study molecule (Figure 5). This fragment was the best ionizable fragment, this guaranteed to identify that fragment and use it in the search for similarity with other molecules described in databases. This short fragment and the results of hemolytic assay corroborate with other mass data analysis to identify our sample as TsTX-κ beta.

**Figure 5.**
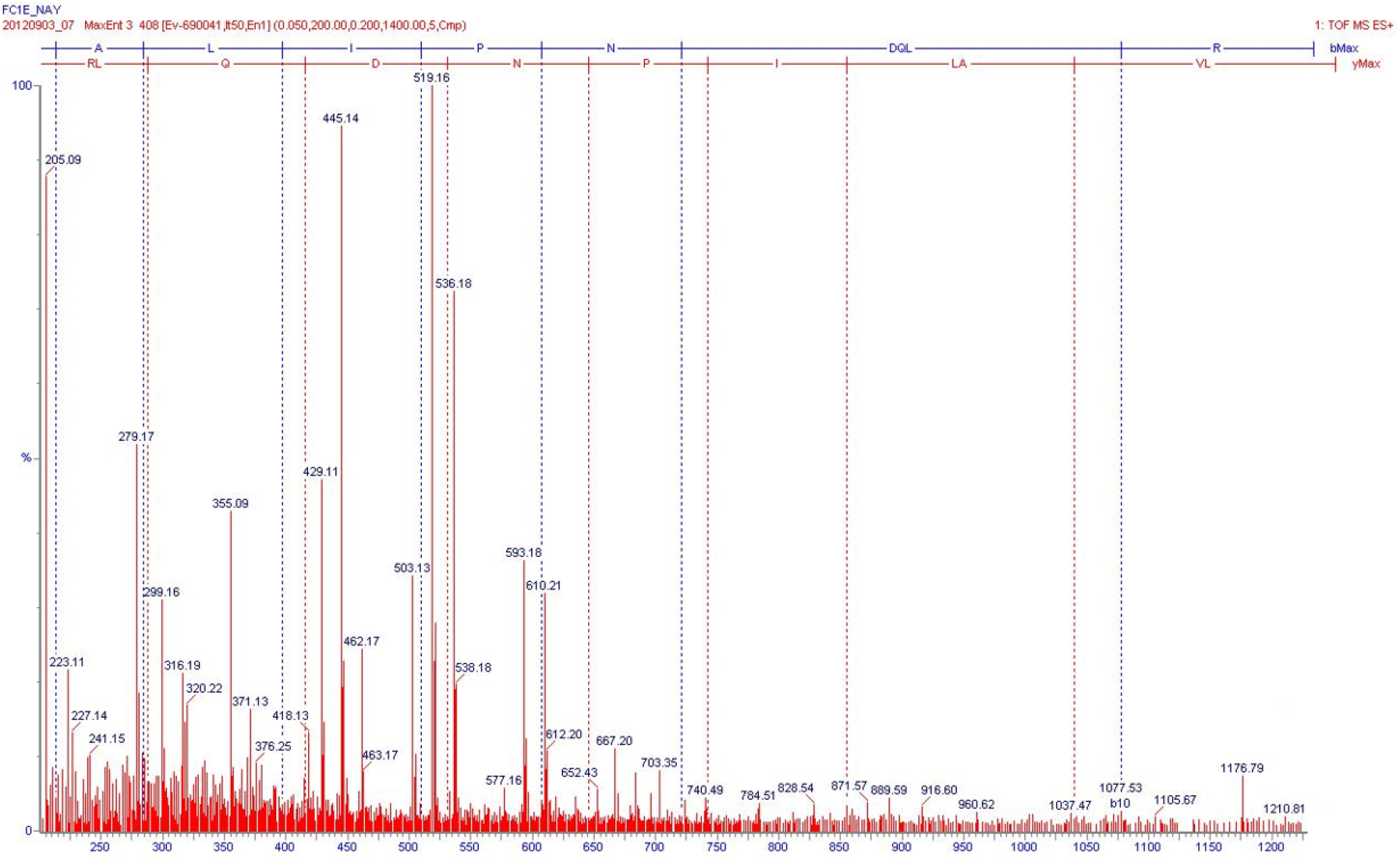
Masslynx sequencing of Fraction 31. The results obtained in the mass spectrometry analysis showed us a sequence of “LVALIPNDQLR” that presents 100% similarity with a neurotoxin with action on the potassium channel called TsTXK-beta from the venom of the scorpion *Tityus serrulatus*.

This same fragment was used in BLAST and UNIPROT, where we can find toxins in other scorpions that share similarity when comparing their primary sequence (Figure 5). Using a short fragment of amino acids identified by mass spectrometry: “LVALIPNDQLR”, it was possible to identify the complete toxin sequence deposited in *T*.*serrulatus* venom transcripts (Uniprot 2025). The complete TsTX-κ beta amino acid sequence was used to search for similarity with other molecules using databases. It is possible to state that similar toxins are found in other species of the same genus, such as *T. costatus; T. trivittatus; T. melici; T. discrepans*. These molecules are toxins conserved in the genus Tityus (Figure 6).

**Figure 6:**
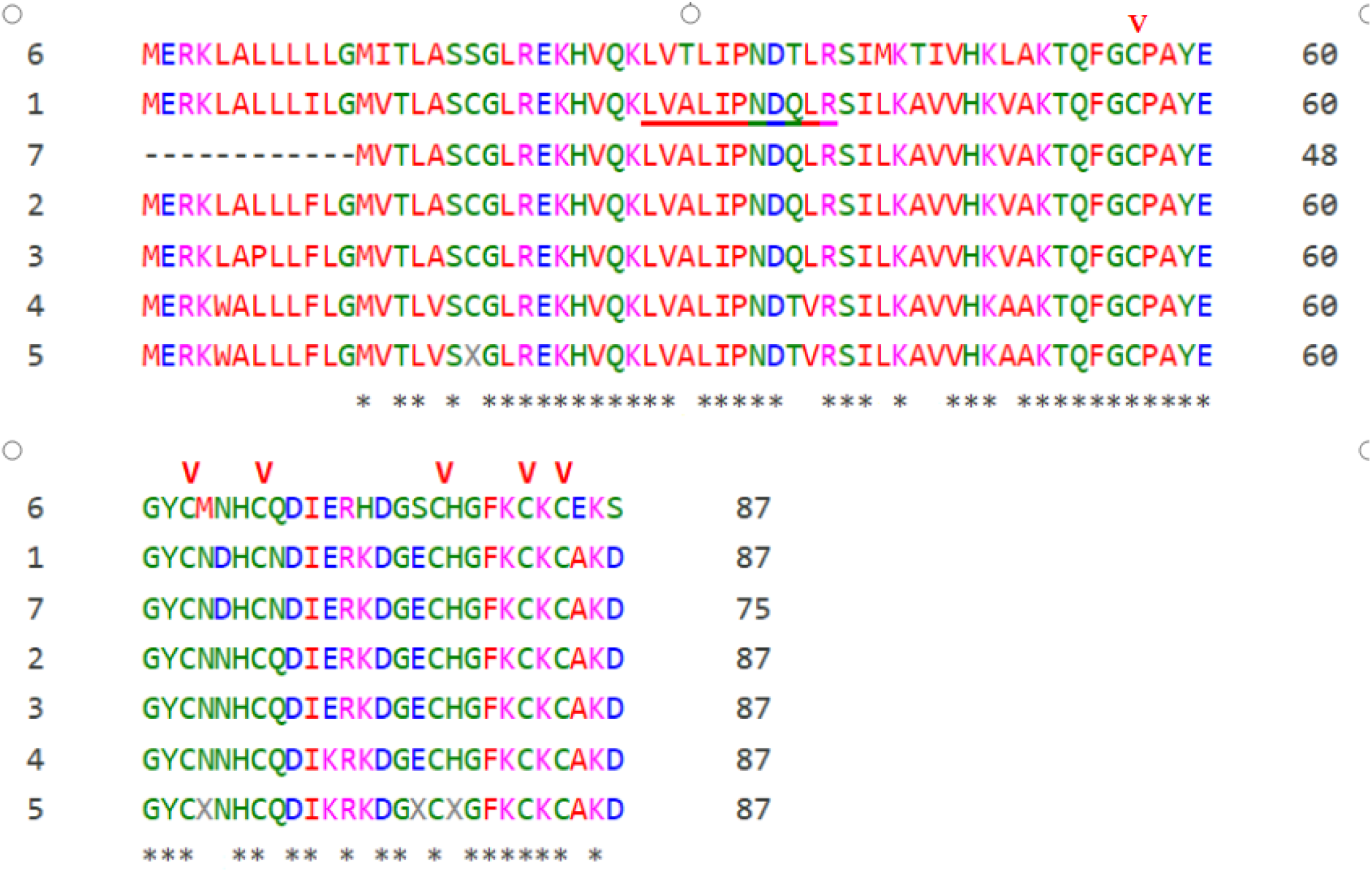
*Alignment analysis of amino acid sequence of TsTX-κ beta from Tityus serrulatus scorpion. This toxin has an amino acid sequence in common with other species of the genus Tityus: 1-T. serrulatus* (Potassium channel toxin TsTXK-beta/Cryptide TyPep-16); 2-*T. costatus* (Scorpine-like peptide Tco 41.46-2); 3-*T. costatus* (scorpine-like peptide precursor); 4-*T. trivittatus* (Potassium channel toxin Ttr-beta-KTx); 5-*T. melici* (putative KTx); 6-*T. trivittatus* (be-ta-KTx-like peptide); 7-*T. discrepans* (Potassium channel toxin Tdi-beta-KTx). Obs.: *(*) the conserved regions and (*V) .*the cysteine residues that form the three disulfide bridges of theses toxin. Underlined, the amino acid sequence that we identified by mass spectrometry and that we used to obtain the complete sequence in transcripts of the T. serrulatus venom*.

Alignment of TsTX-κ beta by Universal Protein Resource (UniProt), indicates that cystine residues form bridges with the following conformation: C1-C4 (56-77), C2-C5 (63-82, C3-C6 (67-84) (UniProt 2025). The position of cystine residues is very common in scorpion venom peptides and is associated with different functions (Gao et al. 2013). This conformation is called inhibitor cystine knot (ICK), and guarantees characteristics such as high chemical and biological stability for the molecule, which are typically resistant to extreme pH, organic solvents and high temperatures (Colgrave and Craik 2004). This molecular conformation is interesting because many antimicrobial peptides from natural sources are not used as antibiotics because they are susceptible to proteolytic degradation. Study with spider venom showed one antimicrobial peptide with ICK motif (Ayrosa et al 2012).

Several antimicrobial peptides were identified in invertebrates in the most diverse sources, such as arachnids, insects and centipedes. In *Loxosceles gaucho* spider venom, the molecule U1-SCRTX-Lg1a was characterized, which acts against Gram-negative bacteria: *Escherichia coli* SBS363, *Escherichia coli* D31, *Pseudomonas aeruginosa* ATCC 27853, *Pseudomonas aeruginosa* PA14 and *Enterobacter cloacae* β-12 (Segura-Ramírez and Silva Júnior 2018).

In spider hemolymphs, it is possible to highlight Gomesin and Rondonin, isolated from the species *Acanthoscurria gomesiana* and *Acanthoscurria rondoniae* respectively (Silva *et al*. 2000, Riciluca *et al*. 2012). Serrulin, a bioactive peptide with antimicrobial activity against Gram-positive and Gram-negative bacteria, filamentous fungi and yeasts was characterized in hemolymph of scorpions *Tityus serrulatus* (Oliveira *et al*. 2019). In addition to the work with arachnids, molecules were identified in other species such as insects, as is the case of *Triatoma infestans*, where antimicrobial molecules that are fragments of human fibrinogen were found in the hemolymph of this insect (Diniz *et al*. 2018) We can also mention the antimicrobial properties identified in the excretion and secretion of larvae of the species *Sarconesiopsis magellanica*, where the action of Sarconesin was described (Díaz-Roa *et al*. 2019). In centipedes, of the species *Scolopendra viridicornis*, lacrain, an antimicrobial peptide obtained from the body extract of this species, was isolated and characterized, which acts against different species of Gram-negative bacteria (Chaparro and da Silva 2016).

It is of great value to emphasize the importance of research in search of new candidates for biopharmaceuticals. The sources of bioactive molecules are present in the most diverse environments, and can be found in microorganisms, plants and animals. Furthermore, this work describes the importance of analyzing the possible functions of molecules with already known characteristics, in order to describe their possible multifunctionalities.

## 3. Conclusions

This work identifies an additional function to TsTX-κ beta, toxin of the Brazilian scorpion venom *Tityus serrulatus*. This neurotoxin has antimicrobial activity against different microorganisms and hemolytic activity against human erythrocytes, the hemolytic activity observed in high concentrations. It was also possible to identify the homology of this neurotoxin with other molecules described in other scorpion species.

## 4. Materials and Methods

### 5.1 Animals

*Tityus serrulatus* scorpions were collected in São Paulo and Minas Gerais states (Brazil), under the Permanente Zoological Material Licence n°11024-3-IBAMA and Special Authorization for Access to Genetic Patrimony n°010345/2014-0. They were kept alive in the biotherium of the Laboratory of Applied Toxinology at Butantan Institute (São Paulo, Brazil) and were fed with cockroaches every 15 days, with water ad libitum. The room temperature was maintened between 24 and 26°C.

This research agrees with Ethical Principles in Animal Research adopted by Ethic Committee in the Use of Animals of Butantan Institute and was approved in the meeting of day April 15, 2013 (protocol n°I-1045/13).

### 5.2 Venom milking

The crude venom of *T. serrulatus* was collected by electrostimulation, based on the method described by Lowe and Farrell, 2011 (Lowe and Farrell 2011). An electric shock generator (model AVS-100 - AVS Projetos Especiais, São Paulo, Brazil) was used to apply brief electric shocks (20 V, 10 Hz for 0.5 ms) repeatedly to telson until the scorpion released the venom, which was collected into a 0.5 mL Eppendorf^®^ tube and remained on ice throughout the extraction period. Then it was lyophilized and stored at -80°C.

### 5.3 Reverse-phase High-performance Liquid Chromatography (RP-HPLC)

In order to obtain the antimicrobial molecules, 4 mg of the lyophilized venom was diluted in 2 mL of acidified ultrapure water [0.05% trifluoroacetic acid (TFA)] and centrifuged at 14,000 xg for 5 min. The supernatant was collected and subjected to Reverse-phase High-performance Liquid Chromatography (RP-HPLC) on a Prominence LC-20A system (Shimadzu, Kyoto, KY, Japan), carried out with an analytic column Jupiter^®^ C18 (10 μm, 300 Å, 250 x 4.60 mm) (Phenomenex International, Torrance, CA, USA). Two eluents were used: phase A (0.05% TFA) and phase B [acetonitrile (ACN)/0.05% TFA]. The linear gradient applied was 2% to 80% of phase B, at a flow rate of 1 mL/min over 60 min., ultraviolet (UV) absorbance of the effluent was monitored at 225 nm and 280 nm and the eluted peaks fractions were manually collected, dried in a refrigerated vacuum centrifuge, reconstituted in 500 μL of ultrapure water and stored at -20°C until use.

The fraction with antimicrobial activity was further purified using the same RP-HPLC system and analytic column, but with an elution linear gradient of 20 to 50% of phase B, for 60 min at a flow rate of 1 mL/min. The fractions corresponding to peaks observed in the chromatogram were collected manually, dried in a refrigerated vacuum centrifuge, reconstituted in 300 μL of ultrapure water and stored at -20°C until the use.

### 5.4 Bioassays

#### 5.4.1 Liquid Growth Inhibition Assay

The fractions obtained by RP-HPLC were evaluated for antimicrobial effect by liquid growth inhibition assay *against Gram-negative bacteria: Escherichia coli SBS363, Gram-positive bacteria: Micrococcus luteus A270, yeast: Candida albicans* MDM8 and filamentous fungus: *Aspergillus niger*. Bacteria were cultured in poor nutrient broth (PB) (1 g peptone in 100 mL of water containing 86 mM NaCl at pH 7.4; 217 mOsm).The mid-log phase cultures were diluted to a final concentration of 10^3^ CFU/mL. Fungus and yeast were cultured in poor potato dextrose broth (1/2-strength PDB) (1.2 g potato dextrose in 100 mL of water at pH 5.0; 79 mOsm). The mid-log phase cultures were diluted to a final concentration of 10^5^ CFU/mL.

The assay was performed using a five-fold microtiter broth dilution in 96-well sterile plates at a final volume of 100 μL. 20 μL of each sample collected by RP-HPLC were applied into the wells and the volume was completed with 80 μL of the microbial culture dilution. 100 μL of sterile water and PB or PDB (the same used in the experiment) were employed as quality controls, they were applied into the side wells (columns 1 and 12). Streptomycin and/or tetracycline were also used as controls of growth inhibition. The microtiter plates were incubated for 18 hours, at 30°C, under constant shaking, growth inhibition was determined by measuring absorbance at 595 nm in a Victor3 microplate reader (Perkin Elmer Inc., Waltham, MA, USA) (Hetru and Bulet 1997, Bulet 2008).

#### 5.4.2 Minimum Inhibitory Concentration (MIC)

Determination of minimum inhibitory concentration (MIC) was performed as anteriorly described, but using a serial dilution of the samples. MIC was expressed as the interval [a]–[b], [a] is the highest sample concentration at which the microorganism grows and [b] is the lowest concentration which inhibits completely the growth Ehret-Sabatier *et al*. 1996).

#### 5.4.3 Hemolytic Assay

Type A+ blood was collected from a human donor and preserved in 150 mM sodium citrate buffer (pH 7.4). It was centrifuged for 15 min at 700 xg, plasma was discarded and red cells were washed 3 times in phosphate-buffered saline (PBS) 1X (137 mM NaCl; 2.7 mM KCl; 10 mM Na2HPO4; 1.76 mM KH2PO4; pH 7.4). At each washing new centrifugations were made and supernatant was discarded. The red cells obtained were diluted in the PBS 1X buffer to a final concentration of 3% (v/v).

Aliquots containing the TsTX-κ beta fragments were dried in a vacuum centrifuge and reconstituted in the PBS 1X buffer, then they were tested in serial dilutions (in duplicate). PBS 1X and 0.1% Triton X-100 solution were used as negative and positive hemolysis control, respectively. The test was performed on a 96-well U-bottom microplate. Into each well, which already contained controls and the diluted peptide, 50 μL of the 3% red cell solution were added. The plate was incubated at 37°C for 1 hour and then the supernatant was transferred to a 96-well flat-bottom microplate and hemolysis was assessed by measuring the absorbance at 405 nm, using a Victor3 microplate reader (Perkin Elmer Inc.). The hemolysis percentage was estimated concerning a 100% lysis control (Triton X-100) and the calculation was made according to the equation: % hemolysis = (Absorbance sample - Absorbance negative control)/(Absorbance positive control - Absorbance negative control) (Hao *et al*. 2009).

The use of human blood was authorized by Committee from the University of Sao Paulo School of Medicine (USP) FMUSP, number: 17526619.3.0000.0065.

### 5.5 Sodium Dodecyl Sulfate-Polyacrylamide Gel Electrophoresis (SDS-Page)

The SDS-Page was performed based on the protocol described by Laemmli (1970). 50 μg of TsTX-κ beta in a non-reducing condition, were analyzed by non-reducing conditions, they were diluted in a 4-fold concentrated buffer (250 mM Tris-HCl pH 6.8; 300 mM SDS; 1 mM bromophenol blue; 40% glycerol) and applied into a 20% polyacrylamide gel (DIGEL vertical electrophoresis system; dimensions: 85 x 80 x 1.0 mm). Precision Plus Protein™ All Blue (Bio-Rad, Hercules, CA, USA) was used as a reference for molecular masses.

The run was performed at room temperature at a voltage of 120 V, using a running buffer composed of 250 mM glycine, 25 mM Tris-HCl and 3 mM SDS. The gel was stained with coomassie brilliant blue R-250 (45% methanol; 10% acetic acid; 3 mM coomassie brilliant blue R-250) and destained by immersion in a 25% methanol/8% acetic acid destaining solution until appearance of blue-colored bands.

### 5.6 In-solution Digestion

Aliquots containing TsTX-κ beta peptide were dried in a vacuum centrifuge and reconstituted in 20 μL of 8 M urea/0.4 M ammonium bicarbonate solution. 5 μL of 45 mM dithiothreitol solution (Invitrogen, Carlsbad, CA, USA) were added and incubated for 15 min at 50°C. After cooling at room temperature, 130 μL of ultrapure water and bovine trypsin (Sigma-Aldrich, San Luis, MO, USA) were added in the 1/25 ratio (enzyme/protein). The peptide was incubated for 12 hours at 37°C, then the digestion was stopped with 0.1% TFA (Stone and Williams 2002).

### 5.7 Mass Spectrometry and Bioinformatics Analyses

The sample obtained after in-solution digestion was resuspended in 20 μL of 0.1% formic acid (FA) and analyzed by liquid chromatography coupled with tandem mass spectrometry, using an LTQ XL (Thermo Fisher Scientific Inc., Waltham, MA, USA) connected to Easy-nLC 1000 (Thermo) system. 11 μL were automatically injected into a pre-column Jupiter^®^ C18 (10 μm; 100 μm x 50 mm) (Phenomenex) coupled to an analytical reverse-phase column Aqua^®^ C18 (5 μm; 75 μm x 100 mm) (Phenomenex). Two eluents were employed: phase A (0.1% FA) and phase B (0.1% FA/ACN). It was used a linear gradient of 5 to 95% of phase B over 18 min, with a flow rate of 200 nL/min. Electrospray ionization (ESI) was operated in positive mode, with temperature and voltage set at 200°C and 2.0 kV, respectively. The mass interval considered for full scan (MS1) was 200-1800 m/z, using the data-dependent acquisition mode (DDA). The 5 most intense ions per scan were chosen for collision-induced dissociation (CID) fragmentation (Zhang and Marshall 1998).

The spectrum obtained was processed by MagTran^®^ to assess the molecular weight of the peptide and by PEAKS Studio software, version X Plus (Bioinformatics Solution Inc., Waterloo, ON, Canada). The searches involved 0.5 Da error tolerance for precursor ions and fragment ions. Carbamidomethylation of cysteines was determined as a fixed post-translational modification. Asparagine and glutamine deamidation, methionine oxidation, and N-terminal acetylation were determined as variable modifications, and trypsin was considered enzyme-specific.

## Acknowledgments and Author contributions

All authors contributed substantially to the production of this work.

We would like to thank Nayrob Pereira for acquiring the initial results during her internship in our laboratory. Dr. Katie Cristina Takeuti Riciluca for her incredible co-supervision in the first steps and discussions. MSc. Soraia Maria do Nascimento for helping with venom extraction, keeping the scorpions in our bioterium and participating in the sample purification steps. We would also like to thank Dr. Ursula Castro de Oliveira for all her help in searching and analyzing the transcripts in the *T. serrulatus* database. And finally, thank to Dr. Pedro Ismael da Silva Junior for supervising this work.

